# Protein Structure Determination via Distance Geometry Structural Similarity based Initialization

**DOI:** 10.1101/2020.10.14.339903

**Authors:** Abiy Tasissa, Tiburon Benavides, Rongjie Lai, Chunyu Wang

**Affiliations:** Department of Mathematics, Tufts University; Department of Biological Sciences, Rensselaer Polytechnic Institute; Department of Mathematics, Rensselaer Polytechnic Institute; Department of Biological Sciences, Department of Chemistry and Chemical Biology, Center for Biotechnology and Interdisciplinary Studies, Rensselaer Polytechnic Institute

## Abstract

The problem of finding the configuration of points given partial information on pairwise inter-point distances, the Euclidean distance geometry problem, appears in multiple applications. In this paper, we propose an approach that integrates structural similarity and a nonconvex distance geometry algorithm for the protein structure determination problem. When initialized with a homologous structure, reconstruction of ubiquitin structure with our non convex algorithm resulted in an RMSE of less than 2 Å with 1.5% available inter proton distance and up to 20% relative error in the input distances. To test the robustness of this algorithm with regard to initialization, we also initialized with a nonhomologous structure on a larger protein with pdb coordinate 1W2E. Even though the initialization structure 1JYB is far different from 1W2E with an RMSE of 25 Å, reconstruction generated structures with RMSE of close to 2 Å, using 1.7% available proton distances and up to 10% relative error in input distances. These results suggest EDG-based approach may be applied to fast NMR structure determination in the future.

## 1 Introduction

The Euclidean distance geometry problem, abbreviated EDG hereafter, is the problem of finding the configuration of points given partial information on pairwise inter-point distances. The problem is of central importance in many tasks such as protein structure determination [1, 2, 3, 4], sensor localization [5, 6] and dimensionality reduction [7]. Formally, consider a set of *n* points **Q** = {**q**_1_, **q**_2_, …, **q**_*n*_} ∈ ℛ^*n×r*^ in an *r* dimensional space and with **D** denoting the partially observed squared distance matrix. The exact EDG problem can be formulated as follows

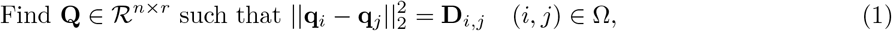

where Ω denotes the set of observed indices. The above minimization problem also has the following compact representation

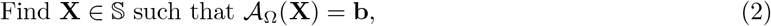

where **X** = **QQ**^*T*^ denotes the symmetric positive semi-definite Gram matrix and the linear operator 𝒜_Ω_(**X**) : ℛ^*n×r*^ →ℛ^| Ω^ | is defined as 𝒜_Ω_(**X**) = {**X**_*i,i*_ + **X**_*j,j*_ − 2**X**_*i,j*_, (*i, j*) ∈ Ω and **b** = {**D**_*i,j*_, (*i, j*) ∈ Ω}. The set 𝕊 is defined as 𝕊 = {**X** ∈ ℛ^*n×n*^ | **X** = **X**^*T*^ & **X · 1** = **0**}. The constraint **X · 1** = **0** fixes translation ambiguity by centering the points at the origin. Since distance measurements are prone to be corrupted by noise, we relax the equality constraint in the above minimization and consider the following optimization program

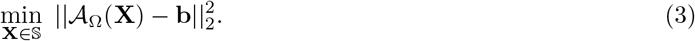

Let **q**^*^ denote the optimal point configuration obtained from solving the optimization program. In this paper, we study the molecular conformation problem where the task is to determine the three-dimensional structure of a molecule given some information about inter-point distances. We focus on the protein structure determination problem where the measurements are obtained from nuclear magnetic resonance (NMR) spectroscopy or X-ray diffraction experiments. Due to the limited range of these measurements, the partial distance matrix is specified in terms of a certain Angstrom threshold. To estimate the protein structure, we make use of a protein with similar structure as the target protein. Hereafter, the point cloud corresponding to the similar protein is denoted by **q**_0_.

## 2 Approach

Consider the exact EDG problem where the *n* atoms in consideration are in a 3-dimensional space with *n* ≫3. It can be verified that the rank of **X** is 3. With this, we can consider a rank regularization on the optimization problem in (2) as follows:

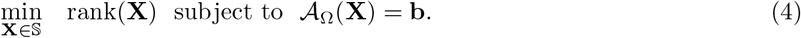

Rank minimization is generally NP-hard [8] and a well known approach is an approximation which replaces the rank function with the convex nuclear norm. The nuclear norm of a matrix is the sum of the singular values of matrix and is denoted by ||**X**||_*_. We apply this convex relaxation to (2) to obtain

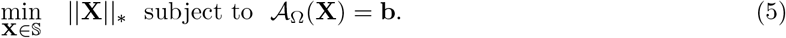

The above problem is a convex optimization program which facilitates ease of computation by employing standard convex solvers. The theoretical analysis in [9], under mild assumptions, shows that solving (5) with *O*(3*n* log *n*) uniformly random distance measurements determines the underlying point configuration with high probability. In practice, the distance measurements are not exact. In addition, the available distance information is not necessarily uniformly at random but rather a deterministic set Ω provided from measurements. In these regimes, the effectiveness of the convex relaxation for the distance geometry problem is not well understood.

One drawback of the program in (5) is its computational complexity. Minimizing the nuclear norm involves the singular value decomposition which can be computationally demanding when *n* is large. Given this, an alternative approach that is amenable to computation is based on factorizing the positive semi-definite Gram matrix **X** as **X** = **QQ**^*T*^ where **Q** ∈ ℛ^*n×r*^. The alternative approach results a nonconvex optimization problem but the minimization over **Q** is generally easier since *r* ≤ *n* in most instances. Given that the nuclear norm minimization is limited to uniform distance measurements, it is of interest to consider a global regularization on the point set that applies in a broader setting. In the context of structure determination, a natural criteria is to enforce that the atoms are at least a certain distance apart. This can be done by considering the minimization of the van der Waals potential term [10]. In particular, we consider the following repulsive force between any pair of atoms [10]

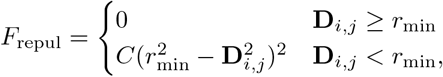

where *C* is a repulsive force constant and 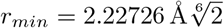. Since the repulsive force is to be minimized over all pairwise distances, this provides a global constraint on the geometry of the points. Another way to enforce a potential term is based on trace of the Gram matrix. We first note the following relation for a centered configuration of point

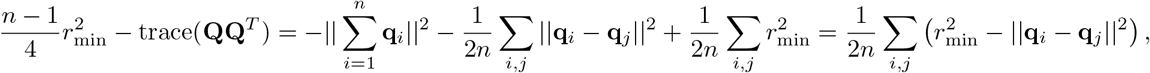

where the second equality uses the assumption that the points are centered at the origin. Hence, akin to the potential term that encourages atoms to be a certain distance apart, maximizing the trace promotes as large distance as between the atoms while obeying the constraint of known distances. Our numerical experiments on protein structure determination indicate that this regularization is effective. Finally, to account for noise, our linear constraint is modified to 𝒜_Ω_(**X**) = **b** + **s** with **s** modeling the effect of noise. Our proposed nonconvex optimization program is

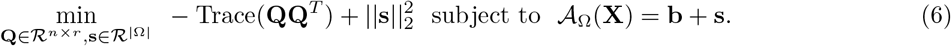

Above, 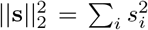 is a noise regularization. Modeling the noise explicitly via this regularization facilities adaptively learning the noise. An advantage is that there is no parameter tuning during numerical experimentation as the noise is optimized as part of the optimization objective. Our numerical experiments indicate that, in the exact EDG setting with *O*(*nr* log *n*) uniformly random measurements, the algorithm corresponding to the above optimization problem typically converges in few iterations to the desired true solution irrespective of the initialization. Compared to its convex counterpart, the algorithm based on the nonconvex formulation is computationally faster. This is due to the employed factorization **X** = **QQ**^*T*^ with **Q** ∈ ℛ^*n×r*^ and *r* = 3. However, in the general noisy EDG problem, numerical experiments indicate that the initialization of the point cloud is crucial. To this end, the point cloud of the structurally similar model can also be used as a good initialization for the nonconvex EDG problem. To solve the program in (6), we use a similar algorithm as in [9] that employs the augmented Lagrangian method and the Barzilai-Borwein (BB) steepest descent method to update **Q**. The augmented Lagrangian is given by

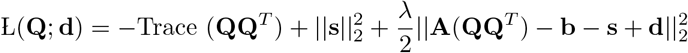

where *λ* denotes the penalty term and **d** is the Lagrangian multiplier. The resulting algorithm, Algorithm 1, is summarized below. **q**^*^ denotes the point configuration obtained from the algorithm. We emphasize that the algorithm is initialized with the point cloud of the structurally similar protein. We posit that this allows us to obtain local convergence for the nonconvex optimization program.

### Algorithm 1 Augmented Lagrangian based scheme to solve (6)

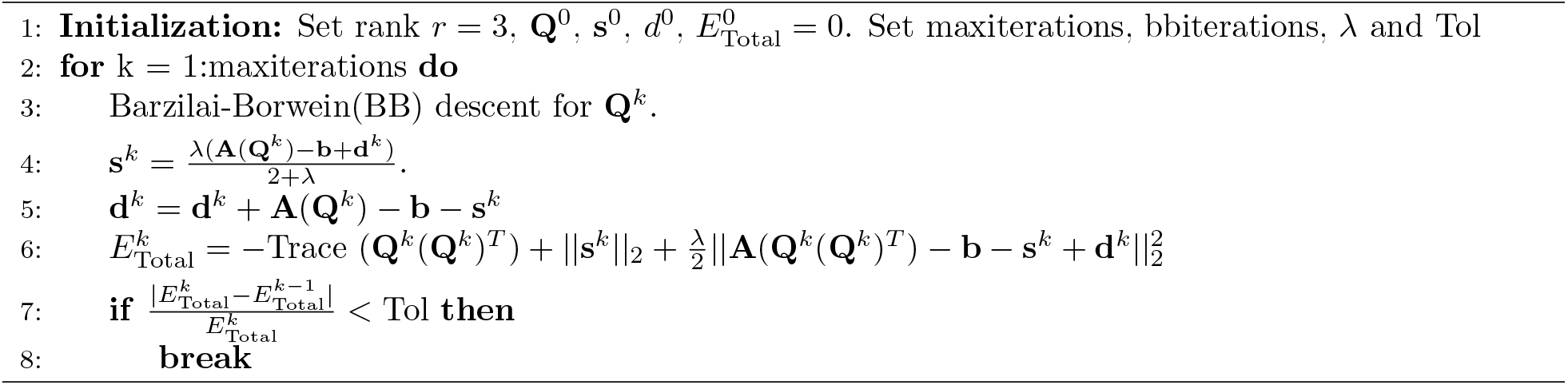

## 3 Results

To test the proposed algorithm, we consider a synthetic experiment to determine the structure of Ubiquitin. This is identified as 1UBQ H, a protonated structure based on 1ubq in the protein data bank and the structure has 629 protons. The structurally similar protein is identified as 2FAZ in the protein data bank. Since pairwise proton distances measured by NMR are not exact, we adopt the following noise model

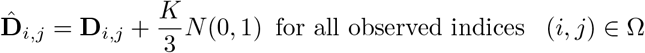

*N* (0, 1) is a Gaussian noise with mean 0 and variance 1 and the parameter *K* controls the range of the noise. The variance is set such that with high probability all the distances are in the range [**D**_*i,j*_ − *K*, **D**_*i,j*_ + *K*]. For our test problem, we first consider a 6 Å cutoff i.e. distances between protons within 6 Å are observed, a range that is typical observed in protein NMR for structure determination. To resemble a realistic setup, from these available distances, we only consider a a subset of the distances i.e. for each point, among the available distances to this point, we pick a certain percentage of randomly selected distances. We initialize the algorithm with the structurally similar model and use the the root mean square error (RMSE) between the numerically estimated point cloud and the ground truth restricted to H-alpha atoms. Table 1 summarizes our numerical results. For small to moderate noise, we obtain good reconstruction of the underlying protein. Given our noise model, there is a trade-off between percentage of available distances and noise level. Since all the available distances are corrupted by a random noise, the input to the nonconvex distance geometry algorithm is more inaccurate when we have more distance samples and a large noise parameter. Fig.1 shows a visualization of the estimated protein obtained from the algorithm. We see a good agreement with the ground truth in the positions of the H-alpha atoms.

**Table 1:** Reconstruction results of 1UBQ H with different noise level (K) and different % of available distances. The parameter *r* denotes the percentage of randomly selected distances among the distances which are within 6 Å for a given atom in consideration. The reconstruction metric is the root mean square error (RMSE) in the unit of Å, restricted to H-alpha atoms, in estimating the structure of 1UBQ H given partial distances. The nonconvex algorithm has been initialized with a homologous protein. The results reported are average of 10 runs.

| noise level | Percentage of available distances |  |  |  |
| --- | --- | --- | --- | --- |
|  | r = 80%<br>(4.26%) | r = 60%<br>(3.20%) | r = 40%<br>(2.10%) | r = 30%<br>(1.53%) |
| K = 0.01 | 0.028 | 0.42 | 0.89 | 1.17 |
| K = 0.05 | 0.033 | 0.38 | 0.97 | 1.25 |
| K = 0.1 | 0.037 | 0.30 | 1.04 | 1.44 |
| K = 0.2 | 0.038 | 0.41 | 1.26 | 1.94 |
| K = 0.5 | 0.067 | 0.54 | 1.71 | 2.96 |

**Figure 1:**
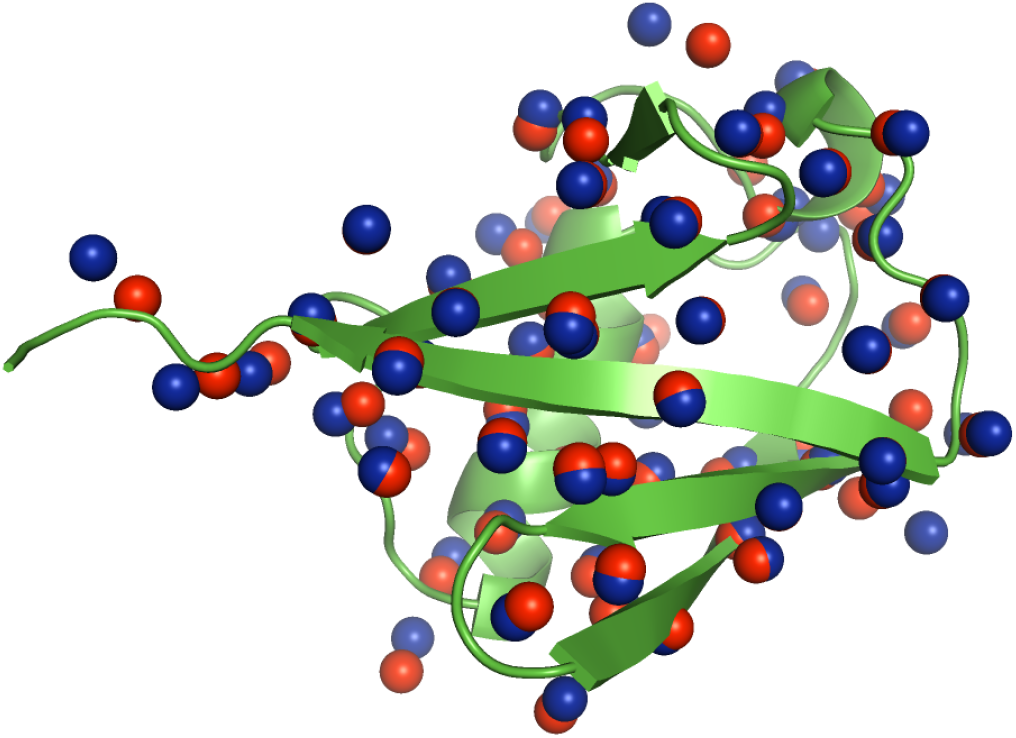
Reconstruction results of 1UBQ H using the proposed nonconvex algorithm. The original structure is shown in green. The red spheres are the H-alpha atoms of 1UBQ H and the blue spheres are the H-alpha atoms of the estimated structure. The algorithm is run with noise parameter *K* = 0.05 and percentage of available distances, among the distances within the 6 Å cutoff, is *r* = 30%.

### Robustness to Initialization

To assess the robustness of the proposed algorithm with respect to an initialization, we consider a synthetic setup in which the initialization is “far” in the RMSE metric from the target protein. The target protein is 1W2E which has 1449 protons. The protein chosen for initialization is 1JYB in the protein data bank, with 1457 atoms. To match the number of protons between the target and initialization model, we select the first 1449 protons uniformly at random from 1JYB as initialization. On average, the H-alpha RMSE between 1JYB and 1W2E is approximately 25 Å. Using the same experimental setup as 1UBQ H, we analyze how the H-alpha RMSE depends on the noise level when we have 80% of the distances within the 6 Å cutoff. Table 2 summarizes our numerical results. Fig.2 shows a visualization of the estimated protein obtained from the algorithm. We see a good agreement with the ground truth in the positions of the H-alpha atoms.

**Table 2:**
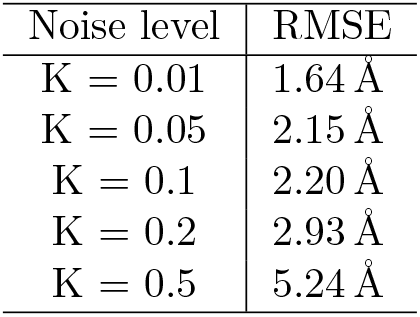
Reconstruction results of 1W2E with different noise levels (K) using 1.17% distance obtained by 80% percentage of randomly selected distances within 6 Å. The reconstruction metric is the root mean square error (RMSE) in the unit of Å, restricted to H-alpha atoms, in estimating the structure of 1W2E given partial distances. To test the robustness of the nonconvex algorithm, it was initialized with 1JYB, which is structurally distinct from 1W2E with an RMSE of 25 Å. The results reported are average of 10 runs.

| Noise level | RMSE |
| --- | --- |
| $K = 0.01$ | 1.64 Å |
| $K = 0.05$ | 2.15 Å |
| $K = 0.1$ | 2.20 Å |
| $K = 0.2$ | 2.93 Å |
| $K = 0.5$ | 5.24 Å |

**Figure 2:**
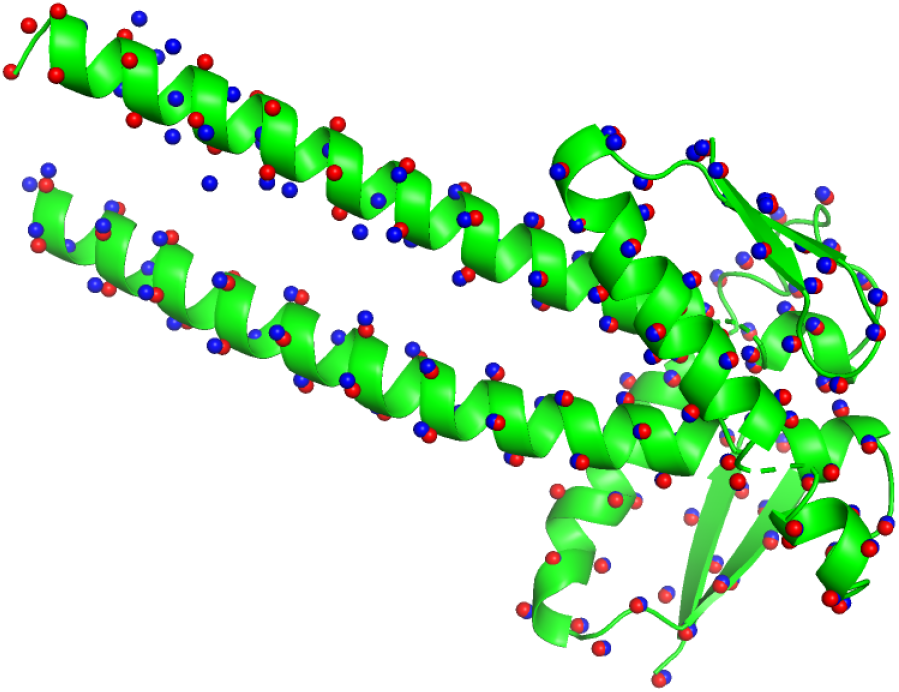
Reconstruction results of 1W2E using the proposed nonconvex algorithm. The original structure is shown in green. The red spheres are the H-alpha atoms of 1W2E and the blue spheres are the H-alpha atoms of the estimated structure. The algorithm is run with noise parameter *K* = 0.1 using 1.17% distance obtained by 80% percentage of randomly selected distances within 6 Å.

## 4 Conclusion

In this paper, we study the protein structure determination problem based on pairwise distances obtained from solution NMR. Using a structural similarity based model, we initialize a nonconvex distance geometry algorithm. Our preliminary results show that, with limited distance measurements and small to moderate noise, reasonable estimates of the underlying structure could be obtained.

## References

[1] Babak Alipanahi, Nathan Krislock, Ali Ghodsi, Henry Wolkowicz, Logan Donaldson, and Ming Li. Determining protein structures from noesy distance constraints by semidefinite programming. Journal of Computational Biology, 20(4):296–310, 2013.

[2] Xingyuan Fang and Kim-Chuan Toh. Using a distributed sdp approach to solve simulated protein molecular conformation problems. In Distance Geometry, pages 351–376. Springer, 2013.

[3] Ngai-Hang Z Leung and Kim-Chuan Toh. An sdp-based divide-and-conquer algorithm for large-scale noisy anchor-free graph realization. SIAM Journal on Scientific Computing, 31(6):4351–4372, 2009.

[4] Michael Souza, Carlile Lavor, Albert Muritiba, and Nelson Maculan. Solving the molecular distance geometry problem with inaccurate distance data. BMC bioinformatics, 14(S9):S7, 2013.

[5] Yichuan Ding, Nathan Krislock, Jiawei Qian, and Henry Wolkowicz. Sensor network localization, euclidean distance matrix completions, and graph realization. Optimization and Engineering, 11(1):45–66, 2010.

[6] Pratik Biswas, Tzu-Chen Lian, Ta-Chung Wang, and Yinyu Ye. Semidefinite programming based algorithms for sensor network localization. ACM Transactions on Sensor Networks (TOSN), 2(2):188–220, 2006.

[7] Joshua B Tenenbaum, Vin De Silva, and John C Langford. A global geometric framework for nonlinear dimensionality reduction. science, 290(5500):2319–2323, 2000.

[8] Benjamin Recht, Maryam Fazel, and Pablo A Parrilo. Guaranteed minimum-rank solutions of linear matrix equations via nuclear norm minimization. SIAM review, 52(3):471–501, 2010.

[9] Abiy Tasissa and Rongjie Lai. Exact reconstruction of euclidean distance geometry problem using low-rank matrix completion. IEEE Transactions on Information Theory, 65(5):3124–3144, 2018.

[10] PJ Kraulis. Protein three-dimensional structure determination and sequence-specific assignment of 13c and 15n-separated noe data. a novel real-space ab initio approach. Journal of molecular biology, 243(4):696–718, 1994.

